# A workflow to gather, clean, and standardize species occurrence records at scale for conservation applications

**DOI:** 10.64898/2026.11.03.763978

**Authors:** Avery E. Pearson, Mitchell Webb, Hannah L. Gray, Shannon M. Collins, Robert L. Whyle, Ross C. Winton, Elinor M. Lichtenberg

## Abstract

Biodiversity declines and pressing threats to wildlife necessitate data-driven approaches for conservation and management planning. Natural resource agencies, conservation practitioners, and researchers increasingly leverage digital species occurrence records. However, we still lack user-friendly and flexible tools to aggregate, clean, and standardize such data. These tools are particularly needed for speciose groups like invertebrates and plants whose natural histories are poorly or unevenly known. To provide such a tool and overcome limitations of existing options, we developed a workflow in R that encompasses the entire process of aggregating, cleaning, and standardizing occurrence records within one free software platform. It additionally can return information useful for assessing species’ conservation status. Key features include user friendliness, flexibility in data sources and filtering, handling large volumes of data or multiple species, and high transparency. We demonstrate workflow functionality focusing on Texas native bees. We combined bee data from the Global Biodiversity Information Facility (GBIF) and several smaller datasets, then cleaned and standardized the combined dataset including removing duplicates. Processing 145,870 bee records took about an hour on a standard-configuration government computer. We used the resulting records in three case studies that demonstrated various uses conservation practitioners commonly implement. Case study 1 compiled a species list for Denton County, the second described phenology of bumble bees (*Bombus* genus) in the Dallas-Fort Worth Metroplex, and the third assessed conservation status of *Bombus pensylvanicus*. Our workflow greatly facilitates addressing a range of questions at various spatial and temporal scales from one vetted dataset that are tailored to the needs of a specific research objective, management question, municipality, or audience.

## Introduction

Reliably identifying at-risk species for conservation prioritization, synthesizing biodiversity trends, and systematically planning conservation requires reviewing the most complete occurrence and natural history data available for one or more species. Indeed, limited or biased data on species occurrences (the Wallacean shortfall), functional traits (the Raunkiærian shortfall), and physiological tolerances (the Hutschinsonian shortfall) can lead to erroneous conservation status or threat estimates, species distribution models, generalizations of ecological concepts, and inferred evolutionary histories (Hortal et al., 2015). Increasingly, natural resource agencies and conservation planners are utilizing occurrence data from online and private databases to more accurately map species’ ranges (Palacio et al., 2021), reduce biases in assessing relative risk among species or land units (Cardoso et al., 2011; Robinson et al., 2016), and plan conservation actions (Cardoso et al., 2011). Occurrence data are used to characterize species’ past and current ranges (Chowdhury et al., 2021; Robinson et al., 2016); estimate physiological tolerances (Hortal et al., 2015); assess species’ conservation status ranks (Bates et al., 2025); determine a region’s or ecosystem’s conservation value (Orr et al., 2022); identify and map surrogate species for conservation planning (Margules and Pressey, 2000); identify under-sampled species (e.g., those ranked as Data Deficient, Bachman et al., 2019), regions, or ecosystems (Hughes et al., 2021) that should be targets for additional research; draft species checklists (e.g., Decker et al., 2020); aggregate community science (e.g., iNaturalist) one-off observations; and track down museum specimens for a given taxon and locale. Achieving these aims often requires combining high-quality datasets from multiple sources (Orr et al., 2022). While data are rapidly becoming more available (Wüest et al., 2020), the lack of user-friendly yet flexible tools to aggregate, clean, and standardize data (IPBES, 2016) slows the application of data-driven approaches in conservation (Cazalis et al., 2022).

Such tools are particularly needed for speciose groups such as invertebrates and plants where data challenges limit natural resource managers’ ability to nimbly generate well vetted datasets to support conservation activities. Data challenges include large numbers of species, sparse data, limited online repositories (unlike, e.g., eBird, Sullivan et al., 2014), lack of standard taxonomic authorities, and effort required to track species’ name changes (Schlesinger et al., 2023). Indeed, while 84% of described vertebrate species have been evaluated under the International Union for Conservation of Nature (IUCN) Red List of Threatened Species, only 2% of invertebrates, 18% of plants, and <1% of fungi and protists have been ranked (IUCN, 2025). Ranking conservation status via the broadly accepted IUCN (IUCN Standards and Petitions Committee, 2024) or NatureServe (Faber-Langendoen et al., 2012; NatureServe, 2020) methodologies requires, at a minimum, an assessment of species rarity based on known occurrences. However, many taxa lack a central repository of observations such as eBird. Wildlife agencies do not consistently maintain the authority and charge to conserve and manage invertebrate and plant wildlife species (Cardoso et al., 2011), and rarely have adequate staffing to cover these diverse groups. For example, a staff biologist may be charged with overseeing conservation of all invertebrate species within a state. Yet, graduate level training usually focuses on a much narrower group of organisms such as arthropods in Entomology programs or freshwater snails in an Ecology program. Additionally, invertebrate conservation biologists will likely have lower levels of support from field biologists within their institution given limited exposure to invertebrates in wildlife degree programs (Grodsky et al., 2015). This often results in limited surveying effort by any one entity and elevates the need to develop workflows capable of combining data from multiple sources to reflect our best knowledge about a given species.

It is often not possible to develop rigorous conservation assessments and plans for at-risk species through publicly available online datasets alone. In an ideal world, all occurrence data would reside in a single database such as the Global Biodiversity Information Facility (GBIF; Orr et al., 2022). In reality, such a database will not include recent, proprietary, or sensitive data. Technical and financial limitations can prevent collections from being made available through GBIF, even if they are online through a different platform (e.g., Alfeus et al., 2025) or as supplemental material for published articles (Mandeville et al., 2021). Thus, relying on public aggregated databases can skew analyses. As Robinson et al. highlighted in their study of aquatic insects, no one dataset is truly complete (2016); they added new state records and new species from a well-studied region. Further, many occurrence records exist in researchers’ “phantom databases” due to plans for publishing recent survey efforts or lack of a standard for data use (Marinho and Beech, 2020). Researchers may, however, be willing to contribute to projects that enable them to keep data proprietary. Additionally, state law may require explicit landowner permission to share surveys on private land publicly, or else those datasets can only be integrated in conservation planning locally at wildlife agencies (Legislature of the State of Texas and Averitt, 2005). Species of conservation concern can be at risk for poaching, and therefore archiving location data publicly is unwise (Feng et al., 2019; Gavin et al., 2010). Given these limitations, natural resource managers need to expand records used in conservation planning to all those available at an ecologically relevant spatial scale (Orr et al., 2022; Robinson et al., 2016) or else run the risk of incorrectly classifying a species ‘rarity’. Such datasets can help overcome geographic and taxonomic sampling biases, particularly in regions where most land is privately held, and for regions and taxa that are under-represented in GBIF (Cardoso et al., 2011; Hochkirch et al., 2021; Hortal et al., 2015). Curation of these data can be very time intensive when completed manually and species by species.

An important next step after exhaustive data source identification is to assemble these multiple datasets into an aggregated, cleaned dataset. The status quo of manual data collation and cleaning (Cornford et al., 2022; Schlesinger et al., 2023) can be prohibitively time consuming especially when assembling data for a large number of species (Bachman et al., 2019). Manual collation can be biased towards the taxonomic interest of the collator, such as an individual researcher or IUCN Specialist Group, or species that are expected to be threatened (Bachman et al., 2019). Natural resource biologists and researchers have largely relied on generating CSV files of observations for a given species and manually inspecting for duplicates or incomplete observations before uploading these data to a spatial program such as ArcGIS or Wallace (Kass et al., 2023) to remove species outliers through visual inspection (e.g., Chesshire et al., 2023; Palacio et al., 2021). Manual removal of observations often occurs without documenting why each observation was removed. This lack of workflow standards limits our understanding of how frequently different types of data errors occur. Relying upon licensed software such as ArcGIS to examine datasets requires institutional investment in software and technical training. Recently NatureServe developed RARECAT, a tool that allows one to pull in GBIF observations as well as a cleaned datafile into an online web portal for visual inspection (NatureServe, 2026). While this is a huge improvement over past tools, the program often crashes with large datasets and in “multi-species” mode. Cleaning a dataset in a standardized workflow would allow for fewer software issues and minimize the number of spurious points that need to be inspected before proceeding with any assessment.

Time and technical expertise limitations also slow species conservation statuses assessment. Several tools have been developed to calculate two spatial metrics important for conducting assessment via the Red List process or NatureServe: extent of occurrence (EOO) and area of occupancy (AOO). These include the browser-based RARECAT (NatureServe, 2026), GeoCAT (Bachman et al., 2011) and sRedList (Cazalis et al., 2024); and R packages ConR (Dauby et al., 2017), redlistr (Lee et al., 2019), and rCAT (Moat, 2020). However, these tools are not sufficient to provide a flexible workflow that can assess species at scale. ConR and rCAT are no longer easily available and do not appear to be maintained. GeoCAT and RedList are limited to providing information for one species at a time (Bates et al., 2025). They also limit the number of occurrence records that can be uploaded (e.g., 5,000 currently for RARECAT), and do not provide a clean workflow for combining data from aggregated datasets and other sources. redlistr functions also assess one species at a time, although can easily be applied to multiple species within the R environment. However, it relies on spatial data such as raster images, rather than simpler lists of species occurrences. Using existing tools requires combining multiple tools for data assembly, cleaning, and calculation (as in Bates et al., 2025). Additionally, most tool development focuses on the IUCN Red List. Many US states, however, use NatureServe. Combining the entire process in one workflow that is user friendly and does not require software licenses is of utmost importance to efficiently and accurately establish baseline species conservation statuses when personnel are limited and biodiversity is high (Hochkirch et al., 2021).

To overcome these limitations, we developed a user-friendly, flexible workflow in R (R Core Team, 2025) that aggregates, cleans, and standardizes data. This workflow encompasses the entire process of assembling a list of occurrence records within one free and open-source software platform. It additionally can return information that is useful for assessing conservation status including species’ extent of occurrence and area of occupancy, and lists of years, counties, and ecoregions where the species is found. The workflow is designed to allow flexibility in aspects such as which data sources are used and how the data are subsequently filtered. It is written with the advanced beginner R user in mind, and provides extensive instructions for user friendliness and high transparency (as recommended in Cazalis et al., 2022).

We demonstrate workflow functionality with three case studies using Texas native bees: compiling a species list for a given geographic location (Denton County), investigating species’ phenologies (bumble bees), and assessing conservation status (*Bombus pensylvanicus/sonorus*). Texas is the second largest state in the United States. It contains 10 distinct ecoregions that host incredible insect biodiversity and unique plant communities. Texas is home to one of the most diverse bee assemblages in the United States (Ascher and Pickering, 2024; Rousseau et al., 2024), at an estimated 1300 native bee species (J. Neff, pers. comm.). Nationally, native bees are experiencing broad declines in both numbers and diversity with more than 30% of those assessed experiencing some level of imperilment (Cornelisse et al., 2025). With documented species declines and the large number of species, a more automated approach is critical for increasing the rates of conservation assessment and re-visiting older assessments as new data are gathered.

## Methods

### Overview

This workflow automates and streamlines assembly and quality control for large numbers of species occurrence records (Fig. 1). It walks the user through a set of steps that we detail below. First, the user sets up their project space by installing and loading necessary packages, creating a standardized file structure on the local drive, and loading custom scripts that are necessary for this procedure. Second, the user imports or manually enters a list of species. They also import a taxonomic backbone from a local file or from an online source such as GBIF. A taxonomic backbone, or backbone taxonomy, is a database that identifies accepted taxonomic names and maps synonyms to accepted names. Third, the user imports species occurrence records for all taxa on the species list, including accepted names and synonyms. Fourth, the data undergo an extensive cleaning and quality control process. These steps use functions in the BeeBDC R package (Dorey et al., 2023b), which are explained in detail by Dorey et al. (2023a). Throughout this process, the user is prompted to save various intermediate files and snapshots for reviewing their data or restarting the workflow from a previous step. Fifth, the user determines how they want to filter or constrain the data. For example, they might limit data to a specific state or time span. This produces a list of cleaned species occurrence records that can be used for many purposes. Finally, the user has the option to calculate several factors that are typically used to assess conservation status assessment of groups such as invertebrates using NatureServe methodology: number of observations, range extent, and area of occupancy. These factors are calculated separately for each species. Users can also add species-specific information from a spatial data layer such as ecoregions.

**Figure 1:**
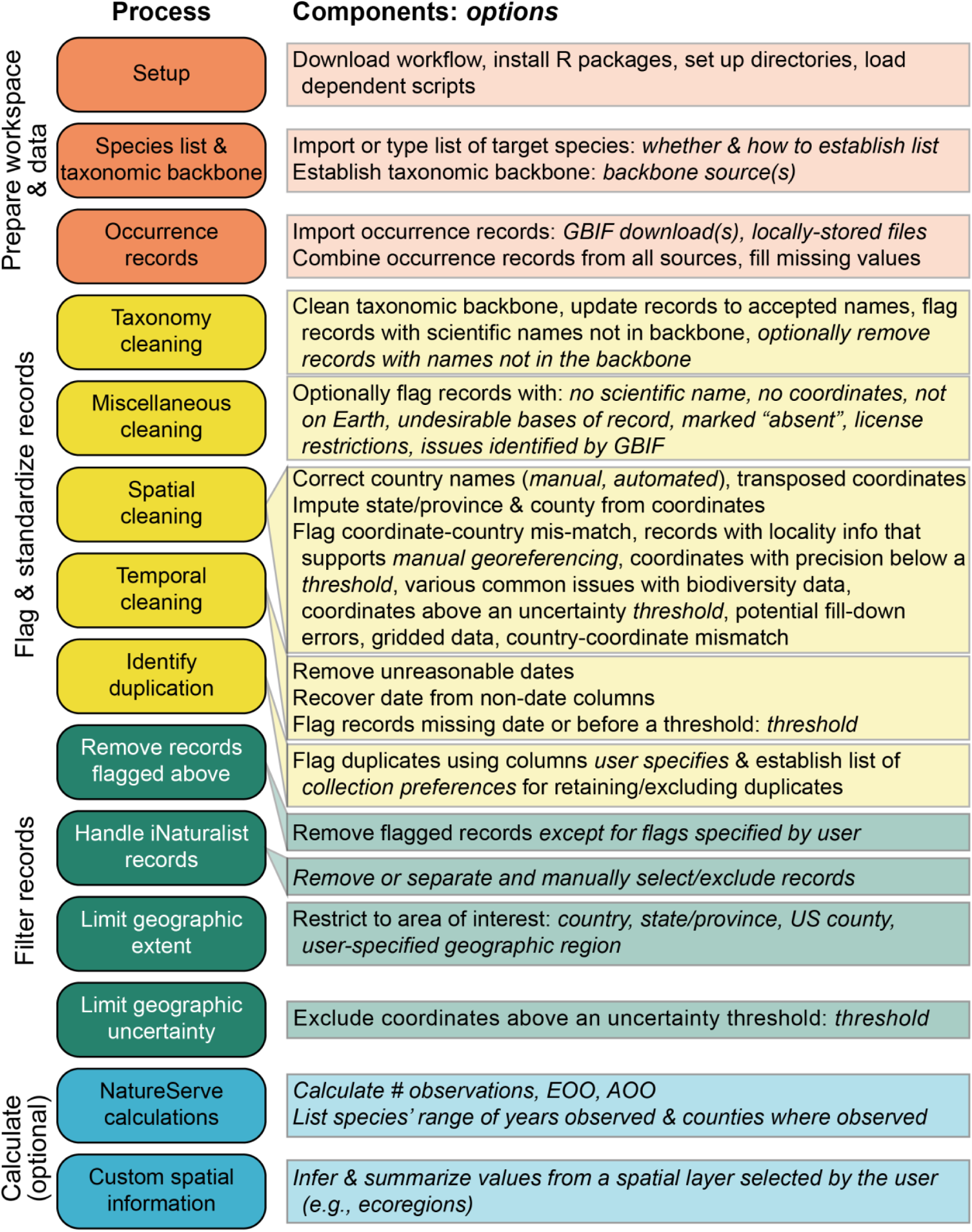
Overview of the steps involved in this workflow, and options available at each step. Italics indicate points at which the user has options.

Our workflow allows customization at multiple steps to best fit the user’s needs and available data. The list of species to obtain records for can be imported from a text file or typed into the script. Advanced users can write code for other options, as well. The taxonomic backbone can be read in from a local file, downloaded from DiscoverLife (bees only), or downloaded via an API from providers such as GBIF. Additionally, the user can generate a backbone that combines several sources to produce a more complete list of species and synonyms. Occurrence records can be obtained from online databases such as GBIF or uploaded locally from proprietary datasets. The script allows for combining data from multiple sources, including downloads from more than one online database. It walks the user through a series of data cleaning steps, many of which can be skipped if they are not relevant (although we recommend implementing most of the steps). Multiple options are provided for filtering the data after data cleaning is complete. This enables our script to be used for a broader range of tasks.

This workflow is available on GitHub (Collins et al., 2026) and can be accessed via direct download/git clone methods by following the instructions in GitHub’s documentation (GitHub, Inc., 2026) or, if Git is installed on the user’s system, through the RStudio New Project dialogue (File -> New Project -> Version Control -> Git). It has been tested using Windows (10, 11) and macOS (Sequoia 15.2). We recommend running it on a computer that has at least 16 GB, and ideally 32 GB, of RAM. It will run with lower amounts of RAM with smaller datasets, but speed may be greatly affected. Prior to running the script, you should install R, RStudio (Posit team, 2026), and GNParser (Mozzherin et al., 2026). R package dependencies are handled at the top of the script. This workflow uses the following R packages: bdc (Ribeiro et al., 2026), BeeBDC, ComplexHeatmap (Gu, 2022; Gu et al., 2016), CoordinateCleaner (Zizka et al., 2019), magrittr (Bache and Wickham, 2025), purrr (Wickham and Henry, 2026), raster (Hijmans, 2025), red (Cardoso and Branco, 2025), rgbif (Chamberlain and Boettiger, 2017), rgnparser (Chamberlain et al., 2023), ritis (Chamberlain, 2021), rnaturalearth (Massicotte and South, 2026), rnaturalearthhires (South et al., 2025), spocc (Owens et al., 2025), sf (Pebesma, 2018; Pebesma and Bivand, 2023), stringi (Gagolewski, 2022), tidyselect (Henry and Wickham, 2024), the tidyverse (Wickham et al., 2019), tigris (Walker and Rudis, 2025), and uuid (Urbanek and Ts’o, 2026). Because the workflow is openly available on GitHub, it can easily incorporate feedback and improvements from diverse users.

### Project setup

To begin using the workflow, the user must first pull or download the entire Invert_occurrence_records GitHub repository. The user then works through the Project Setup section at the top of the Lichtenberg_invert_occurrence_records R script. First, the user sets up necessary R packages. The workflow provides instructions for installing and loading required packages, with sufficient detail to guide someone who is fairly new to using R. After that, the user sets up their file environment. BeeBDC::dirMaker creates a set of directories where species list, taxonomic backbone, occurrence data, cleaned data, and reports will be stored (Fig. 2). Within the /Data directory is a folder named User_data. Any local backbone or occurrence file that the user wishes to use within the workflow should be placed here. The user then runs a line of code to load dependent scripts from the /Scripts directory—the user does not otherwise need to interact with this folder. Finally, the user stores their default graphing parameter values in a hidden variable. The latter may be useful to advanced users who wish to return to their default graphing values following execution of the script.

**Figure 2:**
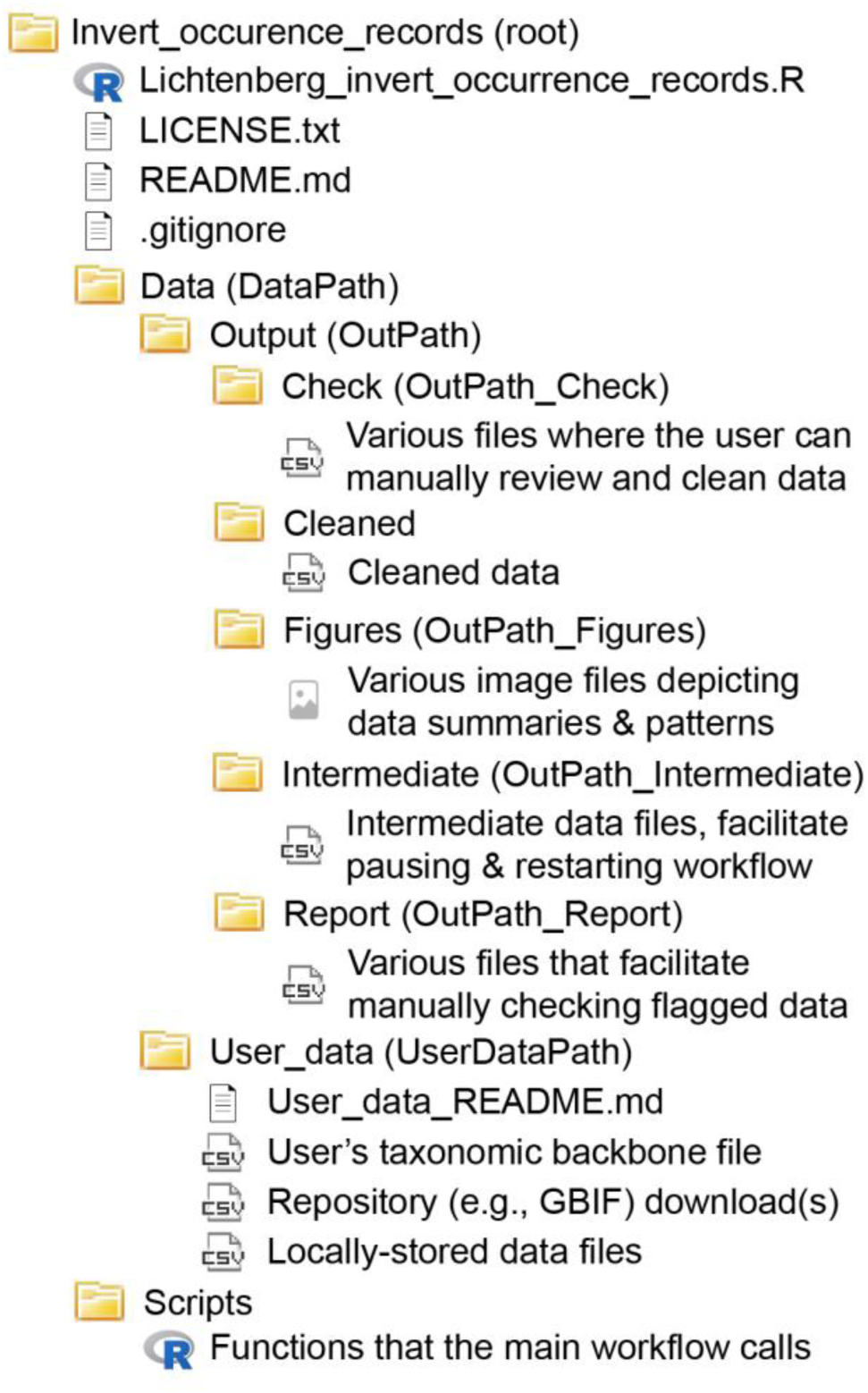
Overview of the files and folders associated with this workflow. The user downloads Lichtenberg_invert_occurrence_records.R, LICENSE.txt, README.md, the Data folder, and the Scripts folder from the GitHub repository. At the time of download, the Data folder will include only the User_data folder and User_data_README.md. All output folders and output files are created while running the workflow. The user places their taxonomic backbone file (if using a locally-stored one), downloads from online databases such at GBIF, and any locally stored data files into the User_data folder. To start using the workflow, the user opens Lichtenberg_invert_occurrence_records.R. LICENSE.txt contains the license for using this script. README.md provides general guidance for using the script, and User_data_README.md describes the User_data file.

### Create species list and taxonomic backbone

The next step in the workflow establishes the species to be investigated and their taxonomic synonyms. Users can optionally specify a list of species to be investigated using the Create Species List section of code. This reduces the amount of data pulled from online databases and improves processing time for data standardization and cleaning. The list can be specified in two ways. First, users can generate a list using an external file with the list of their focal species. Alternatively, users can type their focal species list directly into the script.

Specifying a species list is especially useful when users have one or small group of focal species, like a single genus or species of greatest conservation need (SGCN). However, having a species list is not useful in all situations, like generating a regional species checklist.

The user then specifies their taxonomic backbone. This backbone ensures that scientific names are correct and any outdated synonyms are updated. The taxonomic backbone can be generated from one source or multiple sources. When only using one source, users can read in a taxonomic backbone from a locally-stored or private backbone, DiscoverLife, or an application programming interface (API) like GBIF or the Integrated Taxonomic Information System (ITIS). Locally-stored or private taxonomic backbones are especially well-suited for taxa that have online databases which are outdated or incomplete. DiscoverLife is available only for bee taxonomy. GBIF is well-suited for diverse taxa. ITIS is especially well-suited for plants.

Alternatively, users can combine multiple sources into one taxonomic backbone (following the algorithm in Miller, 2024). This is useful if the most trusted backbone source is incomplete or the focal species list covers a broad range of taxa. To combine multiple backbones, users specify which backbone sources they want to use –locally-stored/private backbones, DiscoverLife, or an API source – and rank each backbone in order of priority from most to least trusted. The backbones are then combined into a master backbone file. When the backbone is later used to verify scientific names, it will use the ranked order. The most trusted source will first be used.

When the most trusted source is missing information for a given taxon, the second most trusted source will be used, and so on. Therefore, the least trusted source is only used when all other sources are missing taxonomic information. Currently, the script is set up to handle up to four backbone sources when creating a multisource backbone, but users can easily add more if necessary. Once the backbone is set up, it will be used to guide the occurrence data queries and data harmonization through the rest of the workflow.

### Import and combine occurrence records

Occurrence records can be imported through an API for online databases such as GBIF or iDigBio, or by importing a local comma-delimited (CSV) file. All datasets are imported into a single object called occDatasets. The workflow currently provides an option for importing data using GBIF’s download API, but the user can add other options with sufficient R coding experience. Importing data from an API can require significant time and processing power, and is recommended only when you are obtaining a limited number of records for a small number of species. When retrieving records from GBIF, the script first compiles a list of GBIF taxon keys associated with the taxa in the user’s backbone. If the user specified a list of species to be investigated, the backbone is limited to those taxa prior to retrieving records. Using the taxon keys list, users can initiate a request for GBIF data using rgbif::occ_download() and download the resulting dataset using get_typed_gbif_download().This section also provides the option to use a previously downloaded GBIF dataset. Note that we use “()” to indicate a function and refer to scripts unique to our workflow (e.g., get_typed_gbif_download()) simply by name and functions from other packages using the format package name::function name.

Users can also import any number of locally-stored datasets. When importing a local dataset, it is critical to consult the list of required columns shown at the top of this section of code. If one or more of these required columns are not included, add and initialize them with NA values when acceptable to do so. Otherwise, the dataset will be rejected with an error displaying missing column names. The data must be in Darwin Core format, the currently accepted standard for biodiversity data (Wieczorek et al., 2012).

Once the user has imported all desired datasets into occDatasets, they are guided to merge all datasets into a single table. This table is formatted one final time to fill missing columns with NA values. From here, the original datasets are removed from the session for processing efficiency. The resulting dataset can be exported to a file for future use.

### Clean and flag data

This section largely uses the existing functions and workflow from the BeeBDC and CoordinateCleaner packages for cleaning and standardizing occurrence data. This includes both harmonization, where established data structures and information such as taxonomic names are applied to the inputted data, and flagging. Flagging steps create a “flag” variable that is set to TRUE or FALSE, depending on the flag, when a record had a potential issue. Comments help the user understand how to interpret flag values, since some are counter-intuitive. This enables users to later filter data as they see fit for manual verification or elimination, as recommended by Orr et al. (Orr et al., 2022). Here we outline the steps involved and indicate differences between our workflow and the workflow outlined by Dorey et al. (Dorey et al., 2023a). Cleaning and standardizing takes place over five main sections: taxonomy, miscellaneous flags, space, time, and duplicates.

The taxonomy section flags and cleans issues related to taxonomic names. In contrast to the BeeBDC workflow, this section occurs near the beginning of our script and added the option to filter occurrence data by desired taxa. Doing so can reduce the number of records and substantially reduce workload in future sections. Taxonomy cleaning first uses bdc::bdc_clean_names() to flag, parse, and clean scientific names in the user’s taxonomic backbone, creating a names_clean column. It then uses BeeBDC::harmoniseR() to flag occurrence records that do not match the cleaned taxonomic backbone. This flag can be used to filter the occurrence data down to relevant records to reduce unnecessary workload in future sections. Filtering with this flag is not required. The user can also use this flag to manually check and correct any occurrence records that do not match the backbone, if they choose.

The miscellany section contains much of the workflow found in the ‘Initial flags’ section of the BeeBDC workflow, although some spatial functions have been moved to future sections. Using various BDC and BeeBDC functions, the user can flag records with empty scientific names, empty or impossible (e.g., not on Earth) coordinates, undesired evidence type (e.g., FossilSpecimen, MachineObservation), occurrence status of “absent”, licenses that indicate the data cannot be used, or issues previously flagged by GBIF. As the user runs the script, BeeBDC::flagRecorder() can be used to collect and view information on flagged records.

The space section contains the workflow found in the Space section of the BeeBDC workflow, some additional state- and county-level imputation functions, and several of the geography-related functions from BeeBDC’s Initial Flags section. Many of these functions use map shapefiles sourced from rnaturalearth. Although the script defaults to using rnaturalearth’s highest-resolution shapefiles, this can still result in inaccuracies on coastlines and near complex borders. The space section first cleans and standardizes country names. With BeeBDC::countryNameCleanR(), the user can manually define country name issues to be corrected. Common examples include inconsistent capitalization or punctuation, such as USA versus Usa or U.S.A. The user is then presented with the option to derive country names from record coordinates, using BeeBDC::jbd_CfC_chunker(). This function is very RAM-intensive, so can take time to run. If the function takes too long, it may be necessary to filter the data or split records into separate objects prior to working through the space section of the workflow. Next, bdc::bdc_country_standardized() creates the countryCode column that contains country codes in a standardized ISO format. This column is used for cleaning coordinates. The user then re-runs BeeBDC::jbd_CfC_chunker() in case more country names can be derived. Now the user is ready to clean and standardize record coordinates. BeeBDC::jbd_Ctrans_chunker() flags and corrects records with coordinates that appear to be transposed. This function is in the Initial Flags section of the BeeBDC workflow. BeeBDC:jbd_coordCountryInconsistent() flags inconsistencies between country and coordinate values, and allows the user to input a buffer value. The prior two functions use the countryCode column. bdc::bdc_coordinates_from_locality() flags issues with potentially useful coordinate information in their “locality” column.

Although sources like GBIF strive to keep data consistent and clean, it is ultimately up to the data provider to enter information correctly. Sometimes key data, like coordinates, can end up in text columns like ‘locality’. The bdc_coordinates_from_locality function can be used to filter the dataset to records that the user may want to georeference. Our workflow provides instructions for extracting these points and adding any corrected information into their dataset. This process must be done manually, as assigning location to the retrieved data may require user insight. The next step in the workflow guides the user through flagging records with undesirable coordinate precision, undesirably large coordinate uncertainty, or various spatial issues. Examples of the latter include coordinates close to universities, zoos, herbaria, country capitals, or GBIF headquarters. This step uses various BeeBDC and CoordinateCleaner functions. Next, BeeBDC::diagonAlley() flags records with sequential coordinates, which may indicate fill-down errors that will need to be manually corrected. CoordinateCleaner::cd_round() flags potentially undesirable datasets with gridded data points. The gridding threshold can be manually set to what is acceptable for the user’s use-case. Although this rasterization may not render data unusable, it may affect the precision and trustworthiness of a dataset. This function can take a long time to run. At this point, the user can compare their occurrence dataset to a user-provided country checklist, outputting a list of failed species-country combinations. The custom checklist must include the following columns: validName as "Scientific name + authorship" (e.g., "Bremus americanorum (Fabricius, 1775)"), iso_a3_eh with three-letter country codes in the ISO 3166 alpha-3 standard (e.g., "LBN", "MEX", "USA"), and continent (e.g., "North America", "Asia", "Europe"). We provide a Discover Life bee checklist as an example in the workflow. It can also be used when working with bee data. BeeBDC::beesChecklist() can be used to download this checklist, which the user can reference when importing or creating their own. Users are then directed through flagging species not found in the checklist as well as species found outside of their designated countries or continents, and viewing a list of failed species-country combinations. This step can be altered to view species flagged with the continentOutlier flag. Now that spatial flagging and harmonization is complete, the user can generate and view spatial errors on a map and a report of flagged spatial issues.

The time section flags, cleans, and recovers values for date-related fields. It includes a few basic functions to visualize and filter erroneous date information. To start, the user is prompted to plot occurrence date information as a histogram and, using some basic numeric filters, set unreasonable year, month, and day values to NA. BeeBDC::dateFindR() is then used to find missing date information in alternate columns (fieldNotes, locationRemarks, and verbatimEventDate) and populate empty date columns where unambiguous date information can be found. After the workflow retrieves dates to the recovered_dates table, the user is prompted to review the new dates, adjust as needed, and merge them with the original dataset. Next, using bdc::bdc_eventDate_empty() and bdc::bdc_year_outOfRange(), the user can flag eventDates that are missing or from before a user-defined year. Time cleaning is complete, and the user can generate and view reports and figures related to time flags.

The final data flagging and cleaning step flags duplicate records. The user is prompted to flag duplicate records by providing BeeBDC::dupeSummary with a set of user-defined columns for comparison. Our workflow follows the dupeSummary help file and defaults to identifying potential duplicates based on records’ latitude, longitude, scientific name, event date, observer, catalog number, and institution. Users can alter which columns are used. After flagging duplicates, the user can generate and plot duplicate summaries to observe and verify duplicate information. Duplicates are simply flagged here, not removed.

### Filter data

We implemented five optional filters that users can implement after cleaning and flagging the data. The user can select which – if any – of the filters are relevant for their purposes. The first filter removes flagged records. We recommend removing all flagged records, but the workflow allows users to specify which flags to filter by, if some of the above flags are not important to them. Second, users can optionally filter out iNaturalist records. iNaturalist publishes to GBIF and thus data downloaded from GBIF are likely to include community science records from iNaturalist. While such records can greatly expand the amount of data available, particularly on private (Dickinson et al., 2010) or urban (Callaghan et al., 2020) lands and over larger geographic areas (Díaz-Calafat et al., 2024), community-identified specimens should be treated with caution (Dickinson et al., 2010; Lowe et al., 2026). Workflow users are provided the option to filter out all iNaturalist records or to manually select the iNaturalist records they trust and want to retain in the final dataset. Instructions in the workflow walk the user through exporting, editing, and re-importing iNaturalist records. The third filter selects records within an area of interest using a spatial layer. Users can restrict their dataset to observations within a given country, state, county, or a geographic region they specify. This can be helpful when generating species checklists for specific regions. Alternatively, users can implement the fourth filter, which filters by the “country”, “stateProvince”, or “county” columns rather than using spatial data. This is useful when occurrence data are missing coordinate information but it can be unreliable given spelling and capitalization differences—especially when examining sub-country regions. We recommend using the (third) spatial filter over this fourth filter when coordinate information is available for records, as this will provide the most accurate location-based results. The fifth and final filter removes records based on coordinate uncertainty. Uncertainty indications are common when latitude and longitude are inferred from written location descriptions, or site coordinates are slightly obscured to protect sensitive species or private land. Users specify a coordinate uncertainty cutoff when querying GBIF. However, any other datasets the user imports are not necessarily filtered by coordinate uncertainty and can be limited here. Once the user completes the filters section of the workflow, their dataset is cleaned, standardized, and ready for analysis. The cleaned dataset is outputted to a CSV file for easy access in the future.

### Calculate values needed by NatureServe

US state wildlife agencies that maintain the authority and charge to conserve and manage wildlife species often use occurrence records to assess conservation status using NatureServe methodology. For taxa like insects where threats and population trends are only generally known, assessment may be based primarily on a species’ rarity. To increase the speed and accuracy with which species can be assessed, our workflow includes an optional function to calculate the rarity factors required for assessment. We include a function that calculates (1) Number of Observations, the total number of occurrence records, (2) Number of Occurrences, unique locations where a species has been observed, (3) Range Extent, the estimated current range of a taxon (also called Extent of Occurrence, EOO), and (4) Area of Occupancy (AOO), the area within the range actually occupied by the taxon (Faber-Langendoen et al., 2012). Number of Occurrences is calculated based on Baker et al (2025). This approach clusters records into “occurrences” based on geographic distance, with a default distance for being a single occurrence of 1 km. Users can change this distance. Our function defaults to calculating distances with sf::st_distance(). Since this function requires significant processing power and time when there are a large number of records, the user may opt to use a slightly less accurate but significantly faster dbscan::dbscan() method by setting the precise_occurrences argument to FALSE. Range Extent is calculated as the minimum convex polygon enclosing all occurrence records, following current best practice (IUCN Standards and Petitions Committee, 2024) and using the sf package. We use IUCN methodology to calculate Area of Occupancy from each occurrence’s latitude and longitude based on a 2 x 2 km grid with red::aoo. Users can optionally calculate the range of years when a species has been observed, and a county list that compiles all counties the taxon is reported to occur in. This function can be applied across a user’s entire dataset to quickly calculate these metrics for each species in even a large list. This will greatly expedite the NatureServe ranking and species occurrence summarizing processes.

### Calculate other spatial information

It can be useful to summarize information related to a species’ distribution other than range and counties where it has been observed. For example, a user might want to know the ecoregion(s) where a species is found. This optional step in our workflow walks the user through determining the categories within a spatial layer where the species is present. The user imports spatial data in a shapefile then intersects the occurrence data with the shapefile. The workflow generates a list of all values for one or more variables of interest within the shapefile where the species occurs. This step requires some experience with shapefiles.

### Case studies

We demonstrated utility of the workflow with several case studies that were designed to simulate common expected uses. Each case study uses a list of Texas native bee occurrence records. To start, we pulled all Texas bee records from GBIF (GBIF.org, 2026a) and read in seven external datasets that were used by Dorey et al. (2023a). We used DiscoverLife for our single source backbone, as this backbone is the most comprehensive and up-to-date source for bee taxonomy. We ran the dataset through the flagging and cleaning portions of the workflow. We originally used the third spatial filter to restrict the dataset to records occurring in Texas but found that the rnaturalearthhires’ (South et al., 2025) Texas boundary was very approximate along the state’s southern border, which follows a river. This led to 1,417 valid Texas records being excluded. We then switched to using a Texas shapefile from the Texas Department of Transportation (Texas Department of Transportation, 2025), which largely avoided this issue. We subsequently corrected or removed flagged records following the workflow’s defaults and records with 2 km or less coordinate uncertainty where measurable. We also excluded iNaturalist records, since there is debate over visual identification of bees and they can require manual curation (Liu et al., 2026).

Case study 1: local species lists. One anticipated common use of this workflow is to generate species inventories that list all species found in a given geographic area (Lowe et al., 2026). We created a bee species list for Denton County, Texas. Using county shapefiles from the tigris package, we used spatial filtering to restrict our cleaned Texas bee occurrence dataset to occurrences in Denton County. We then summarized this dataset using the tidyverse to generate a dataframe listing all the unique bee species and their abundances. Such abundances can indicate which species are locally rare or common.

Case study 2: regional phenology of a genus. Phenological datasets for a given taxon and region are useful when planning surveys to ensure field data collection coincides with the regionally appropriate window of bee activity. We use our cleaned dataset to generate a sampling window for one bee genus within a relevant geographic region based on verified recent occurrences. We filtered the full dataset to contemporary bumble bees, or bees in the genus *Bombus*, that were recorded in the last 30 years: 1995-2025. We also restricted occurrences to those from the Dallas-Fort Worth Metroplex: Dallas, Tarrant, Collin, and Denton Counties. We then grouped the occurrences by species and month of occurrence to identify each species’ first and last months of occurrences and the month with the greatest number of occurrence records.

While abundance values should be used with caution when data come from unstructured sampling, this approach can give a rough estimate of months with higher and lower bee activity that can be useful when structured data are not available.

Case study 3: state-level conservation status assessment metrics. Users can easily use the cleaned occurrence dataset to streamline calculations of rarity using the get_NatureServe_values() function. We calculated rarity metrics for *Bombus pensylvanicus* (sensu lato, *B. pensylvanicus* and *B. sonorus*) in Texas using contemporary records (last 30 years) and generated a list of counties and ecoregions where this species occurs.

## Results

### Workflow performance and Texas dataset metrics

We used the workflow to clean and standardize Texas bee records downloaded from GBIF and from seven external datasets. Once the data were downloaded from GBIF, this process took approximately one hour. A significant portion of this time consisted of several steps that take longer to run: county imputation, transposed coordinate correction (BeeBDC::jbd_Ctrans_chunker), dataset gridding analysis (CoordinateCleaner::cd_round), spatial figure generation (BeeBDC::jbd_create_figures), date recovery (BeeBDC::dateFindR), and the occurrence counting step within get_NatureServe_values (if precise_occurrences is set to TRUE). To provide guidance on necessary processing power and expected run time (Table 1), we re-ran the workflow using randomly sampled subsets of a larger GBIF download that consisted of all 5,380,465 US records in the bee families Andrenidae, Apidae, Colletidae, Halictidae, Megachilidae, and Melittidae (GBIF.org, 2026b). These estimates were made on a Windows laptop with 32GB of RAM. Note that some spatial functions (like get_NatureServe_values) scale exponentially in processing time with the number of records for a given species. With these functions, one species with 10,000 records will take much longer than 5,000 species with 2 records each.

**Table 1:** Estimated processing times for running the workflow on datasets of different sizes. These times exclude data and workflow downloads and package installations.

| **Number of Records** | **Processing Time** | **Suggested Minimum RAM** |
|---|---|---|
| 1,000 | ≈ 10 minutes | 8 GB |
| 10,000 | ≈ 15 minutes | 16 GB |
| 100,000 | ≈ 30 minutes | 16 GB |
| 1,000,000 | ≈ 4 hours (+1.5 hours if using precise_occurrences for get_NatureServe_values) | 32 GB |

Our data pipeline aggregated 145,870 Texas bee occurrence records across all data sources. The cleaned bee occurrence dataset contains 27,2263 occurrences across 815 species. The additional effort of gathering recent survey data that is not in GBIF resulted in a 2.80% increase in occurrence records compared to what was available from public repositories, even with the limited number of local datasets that we used. Of the 118,644 records removed during the cleaning process, 88.67% were iNaturalist observations, 18.59% had coordinate uncertainty > 2 km, 3.70% were duplicates, and 1.00% lacked collection event details.

Our case study highlighted areas where manual data cleaning could improve data quality. First, our workflow does not automate standardization (harmonization) of administrative divisions other than country given that to our knowledge there is no global dataset with all possible permutations of administrative names across languages and spatial scales. This led to a final cleaned dataset where the stateProvince variable included both “Texas” and “texas”. The dataset contained 423 unique county names for the 245 counties in Texas including variations like “County”, “county”, “Co.”, and “co.”; two county names; missing spaces; and no county included in the name. As our case studies depended in part on county assignments, we cleaned county names by coordinating matching with the tigris dataset. This approach cleaned county names for 24.55% of the records, assigned counties to 1.10% of records previously missing any county name, and left < 0.1% of records without a county assignment. Second, our workflow does not automate retrospective georeferencing because current approaches are not sufficiently accurate and can introduce temporal or geopolitical biases (Cornford et al., 2022; Murphey et al., 2004). Inspection of our Texas bee dataset shows that approximately 24,697 records could be added by manually georeferencing records that contain collection location descriptions.

The final dataset contained occurrence records from 1894-2025, with most records occurring during 1950s and 2010s decades (Fig. 3). 99.6% of records were from physical specimens, whereas 0.4% were from ecological occurrence datasets. Our dataset contained records from 226 of the 254 counties of Texas, however only 124 counties had records from the current century (Fig. 4). Occurrences were spread across all 12 of the Level 3 Ecoregions of Texas, with only the Arizona/New Mexico Mountain ecoregion, of which covers less than 0.1% of the entire state lacking contemporary 21^st^ century records. Apidae was the most commonly observed bee family within the dataset, accounting for 36.93% of records, followed by Halictidae and Megachilidae families (Table 2).

**Figure 3:**
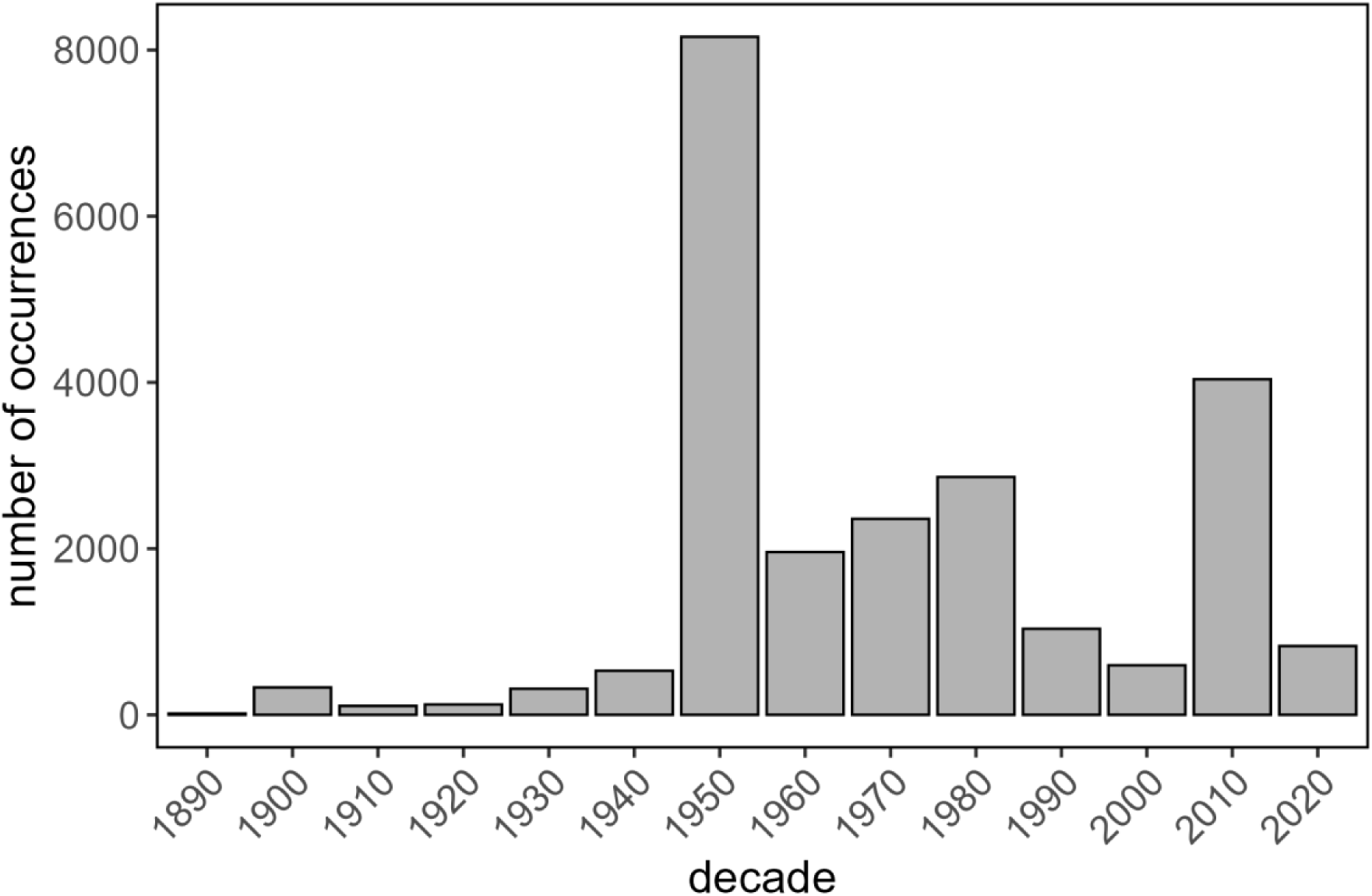
Frequency of Texas bee occurrences by decade.

**Figure 4:**
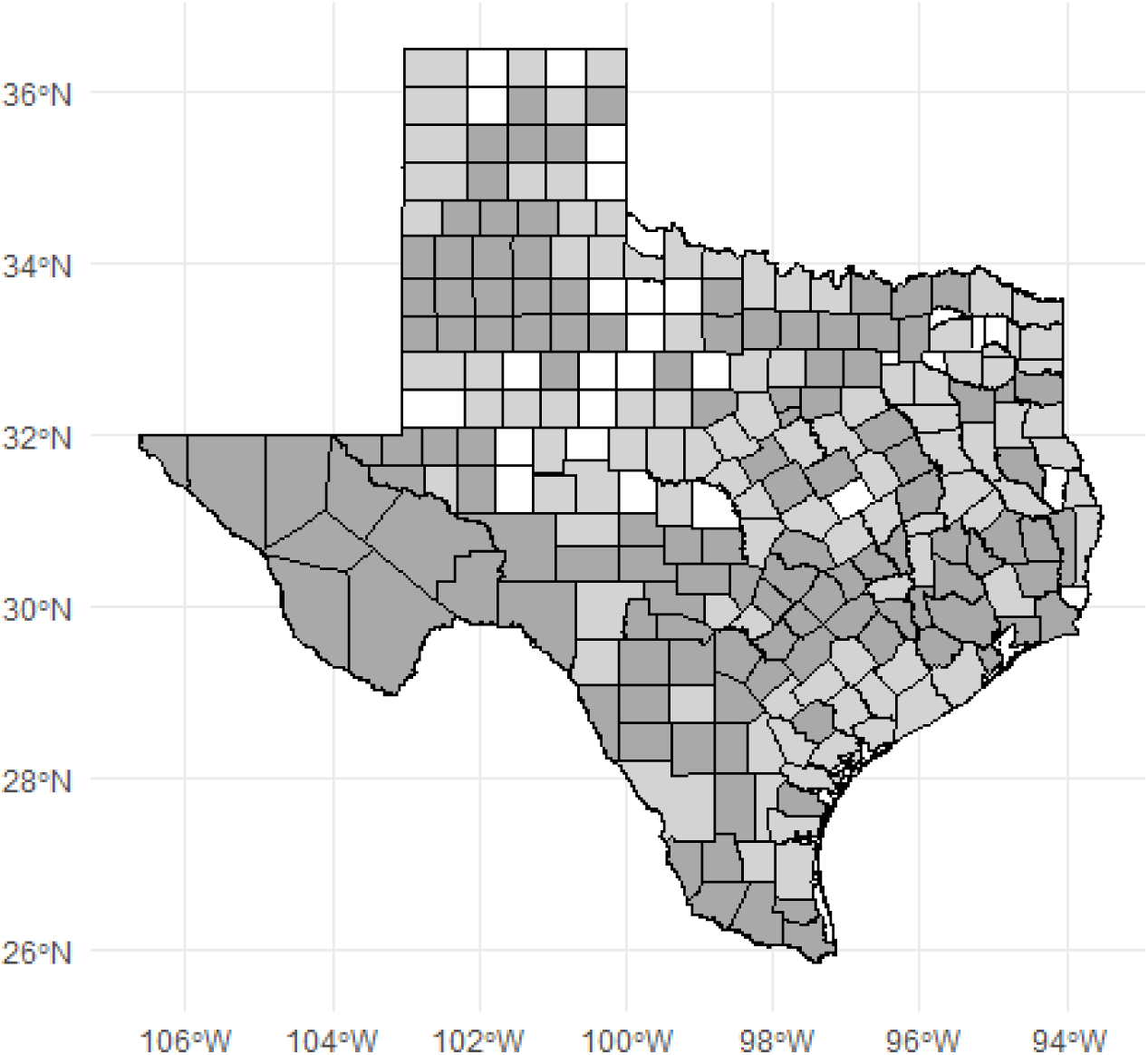
Frequency of bee occurrences by county in Texas. Counties with 21^st^ Century records are shaded in dark gray and those with only historical (before 2000) records are in light gray.

**Table 2:** Year range, number of occurrences, and species richness for each bee family from 1895-2025 in Texas.

| **Family** | **Year Range** | **Number of Occurrences** | **Species Richness** |
|---|---|---|---|
| **Andrenidae** | 1897-2025 | 4930 | 196 |
| **Apidae** | 1894-2025 | 10070 | 259 |
| **Colletidae** | 1900-2025 | 641 | 53 |
| **Halictidae** | 1901-2025 | 6329 | 120 |
| **Megachilidae** | 1895-2025 | 5232 | 183 |
| **Melittidae** | 1949-2005 | 24 | 4 |

### Case study 1: species lists

With our cleaned dataset, compiling a species list for a given geographic location was simple. It took less than a minute to generate a species list based on the 325 occurrences found in Denton County and create a list of the top 10 common bees for this municipality. Our dataset contained 52 distinct bee species in 25 genera and 5 families. The top 10 most abundant native bee species (excluding honey bees, *Apis mellifera*) were *Bombus pensylvanicus* (De Geer, 1773)*, Melissodes boltoniae* (Robertson, 1905)*, Melissodes coreopsis* (Robertson, 1905)*, Anthophorula asteris* (Mitchell, 1962)*, Xylocopa micans* (Lepeletier, 1841)*, Xylocopa virginica* (Linnaeus, 1771)*, Augochloropsis metallica* (Fabricius, 1793)*, Agapostemon texanus* (Cresson, 1872)*, Halictus ligatus* (Say, 1837)*, Lasioglossum disparile* (Cresson, 1872). Data such as these can be used to generate outreach material to municipal departments, local conservation groups, or the general public.

### Case study 2: investigating the regional phenology of a genus

Our cleaned dataset greatly facilitated rapid gathering of phenological data on a single genus within a major metro area. In 1 min we pulled 57 cleaned current records of three *Bombus* species present in the Dallas-Fort Worth Metro Area (Dallas, Tarrant, Collin, and Denton Counites) from 1995-2025 (Fig. 5). The phenological graph in Fig. 5 quickly communicates that most bumble bees in this metro area are *B. pensylvanicus* and that the best window to observe this species is from September to October.

**Figure 5:**
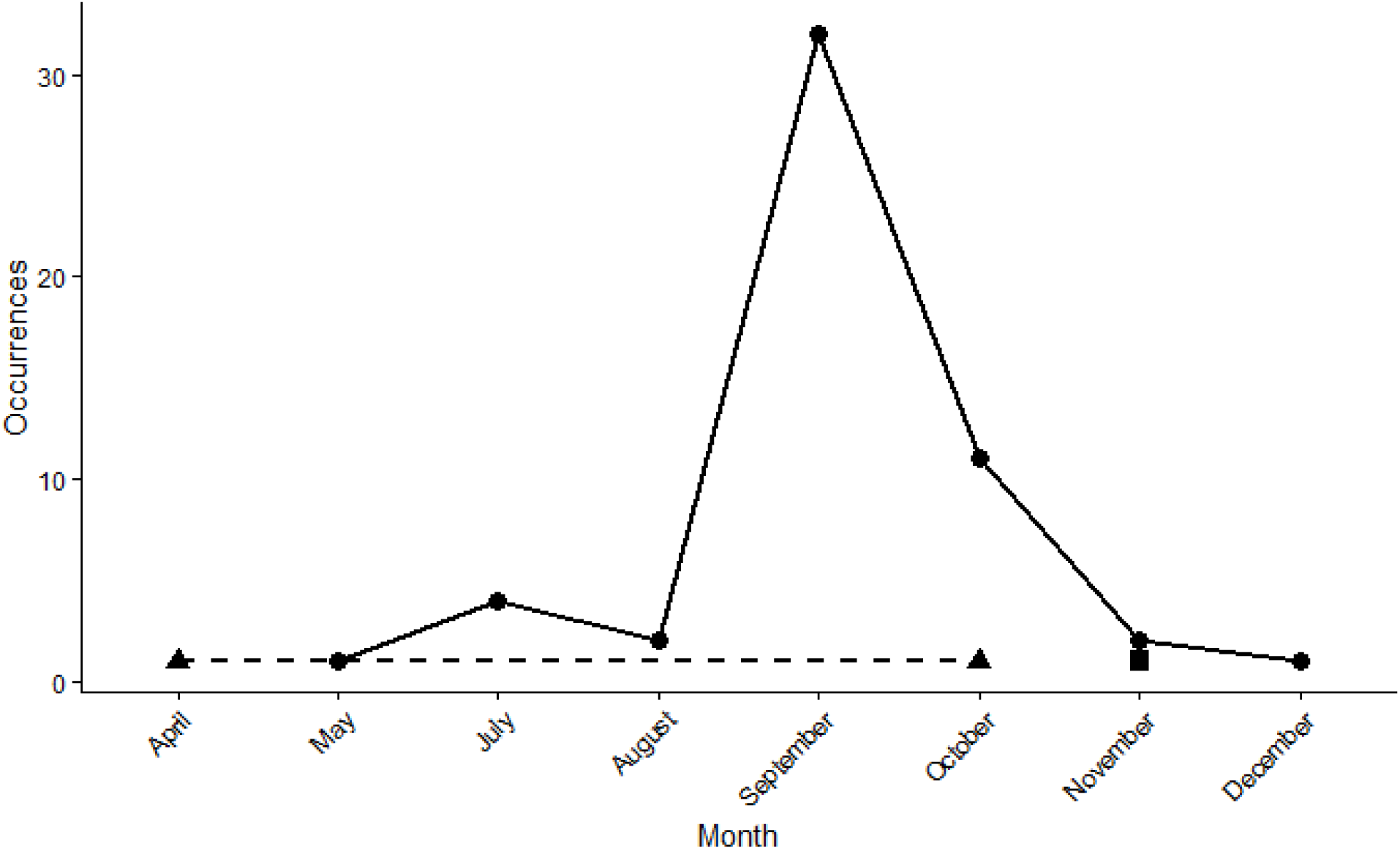
Phenological summary of *Bombus* occurrences in the Dallas-Fort Worth Metro Area from 1995-2025 with *B. pensylvanicus* occurrences depicted with circles and solid lines, *B. fraternus* with triangles and dashed lines, and the sole *B. impatiens* occurrence indicated with a square.

### Case study 3: conservation status assessment metrics

Our dataset quickly generated species spatial metrics needed to initiate a conservation status assessment under both the NatureServe Rank Calculator (NatureServe, 2020) and the IUCN framework. Examining contemporary records of *B. pensylvanicus* (sensu lato including records labeled *B. sonorus*) from the last 30 years (1995-2025), we calculated the range extent as 585,010 km^2^ with 528 km^2^ area of occupancy and 127 occurrences from 361 records located in 69 of the 245 counties in Texas and all 12 Level III Ecoregions. With these minimum spatial rank metrics, the NatureServe Rank calculator assigns the species a preliminary rank of “Apparently Secure” or S4.

## Discussion

Natural resource agencies and conservation planners increasingly rely on digital species’ occurrence records but lack tools to aggregate, clean, and standardize such data at scale. We developed a user-friendly and flexible R workflow as a new tool to improve biologists’ and natural resource managers’ ability to provide data-supported answers to foundational species occurrence questions. The generated vetted dataset enables users to answer species occurrence questions at various spatial and temporal scales, tailored to the needs of a specific management question, municipality, or audience. Our workflow improves on other available tools by (a) handling large volumes of data provided the user has sufficient processing power, (b) facilitating aggregation of multiple datasets including online databases and locally stored proprietary or sensitive data, (c) automating data cleaning, (d) providing information needed to assess conservation status using NatureServe methodology, and (e) handling multiple species all within one tool. It does this with higher flexibility and transparency than some other tools provide. Our workflow is also an improvement over the manual approach that many resource managers currently use. This workflow will be particularly valuable for speciose groups whose natural histories are poorly or unevenly known and thus where natural resource managers rely largely on occurrence records, such as invertebrates and plants (IUCN, 2025).

To illustrate several ways that our workflow can assist with data-driven conservation planning, we used the workflow to assemble a cleaned set of Texas bee records then implemented three case studies. Note that we provide these case studies as examples, but many Texas bee records are not yet databased and thus we do not view these case study results as definitive. Case study 1 showed that Denton County, TX is home to 52 bee species. While the City of Denton became a certified ‘Bee City USA’ affiliate in 2016 (Luxton et al., 2020), publicly available bee outreach material lacks any species-level information. Instead, municipal resources focus on beekeeping practices for non-native European honey bees (*Apis mellifera,* Linnaeus, 1758) and general habitat management for regionally adapted flowering plants (City of Denton, N.D.). A local species list could help the City of Denton switch their Bee City USA activities to providing flowers and shelter targeted to the local native bee species. Case study 2 showed that *Bombus pensylvanicus* is active later in the year than *B. fraternus*. This finding could help local biologists better schedule targeted survey efforts for each bee species or set expectations with regional landowners or public land managers about when they could expect to see active bumble bee pollinators following restoration efforts. Case study 3 assigned *B. pensylvanicus* a preliminary NatureServe rank of "Apparently Secure" within the state, although this species currently petitioned to be listed as federally endangered (Center for Biological Diversity and Bombus Pollinators Association of Law Students, 2021). Such rankings help the Texas Parks and Wildlife Division allocate limited resources for conservation actions.

Our workflow provides several improvements over other available tools. The extensive data standardization and cleaning can increase the utility of individual records by correcting mistakes such as transposed coordinates, and capturing information such as state that could otherwise cause the record to be discarded. Combining automated tasks with flagging of records that are potentially erroneous (as recommended by Orr et al., 2022) or that are targets for additional manual assessment facilitates quality control checking and “recovering” records that can be retrospectively georeferenced or visually inspected. Multiple species and data sources can be processed at once. This is true for both cleaning records and calculating the values needed to assess species with NatureServe methodology (Faber-Langendoen et al., 2012). The latter greatly facilitates bulk assessment of many species with NatureServe’s Rank Calculator (NatureServe, 2020) rather than having to manually enter numbers for one species at a time. Our workflow can also handle large numbers of occurrence records (unlike RARECAT, NatureServe, 2026), limited mainly by available RAM. Our spatial filtering is faster than redlistr (Lee et al., 2019), which uses raster images. With our case study, spatial filtering took just a few minutes. Our workflow completes all steps within one workflow using one free software package. This simplifies the process and makes it accessible to more users. Finally, our workflow allows significant flexibility to meet a user’s specific methodologies and needs. For example, users can integrate data from online databases with local files, including ones that cannot be publicly shared (a current major cause of data gaps, Wetzel et al., 2018). This flexibility was most beneficial in our case study with spatial filtering, when the default method removed Texas data points due to Texas complex southern border but we were able to input a Texas-specific shapefile with higher border accuracy. Users can easily see how applying specific filters, such as using different year cutoffs for data inclusion, alters results. This was most evident in our second case study, where the aim was to determine the current phenology of bumble bees in a metro area. By filtering our results to those in the past 25 years we reduced the number of occurrences by 26% and removed 3 species from our dataset to ensure that results were most relevant for future survey efforts. Our workflow also provided high transparency to easily see how spatial filtering altered which occurrence records were retained with different spatial filters.

Our workflow facilitates several tasks that we did not implement in our case study: incorporating iNaturalist records and retrospectively georeference records without spatial coordinates. These tasks require human intervention (as automated georeferencing currently produces relatively high error rates, Murphey et al., 2004) and can be significant time investments. Our workflow makes it easy to identify and separate such records using the iNaturalist filter and the “coordinates from locality” output file. This automates one step of the process that could otherwise require enough effort to be a barrier. Because our workflow is entirely within one application, users with sufficient R expertise can easily implement additional filters or feed workflow outputs into other analyses. For example, the cleaned dataset can be used for species distribution modeling (as in Doré et al., In review). Batch georeferencing of multiple specimens with the same site description could also be possible with some datasets. Outputs from our workflow can potentially be combined with use of the iNaturalist API to extract additional information about each record and design custom filters.

Our workflow, and any automated processing of species occurrence records, does have some limitations. Occurrence lists and associated range maps and sizes, conservation assessments, or other analyses are limited by available data. Many species have significant data gaps due to limited sampling effort, remote or difficult-to-access habitats, uncertain locality information or outdated identifications for older specimens, or lack of taxonomic expertise and time to identify small and sometimes cryptic species (Bland et al., 2017; Hochkirch et al., 2021). A species’ sampling history can also introduce spatial or temporal biases in available data that can obscure true patterns (Verspagen and Erkens, 2023). Ease of identification can generate taxonomic biases in available data. For example, many butterflies are easily identified with the use of a commercially-available field guide while tiny bees require careful scrutiny under a microscope. Keys and species descriptions do not exist beyond a fairly high taxonomic level for many taxa. While digitization of museum and research collections is accelerating, much of the collected data is not yet broadly accessible (Marinho and Beech, 2020; Stephenson and Stengel, 2020). Indeed, data deficiency is one of the major hurdles to understanding perceived versus true rarity, and conservation needs, of many invertebrates and plants. Data deficiency also may unintentionally direct conservation efforts away from where they are most needed (Bachman et al., 2019; Borgelt et al., 2022). Restrictions on which data are passed to data aggregators like GBIF and on information provided through database APIs additionally restrict outputs from an automated workflow such as ours. iNaturalist, one of the most widely used community science platforms, only exports Research grade observations with CC0, CC BY, or CC-BY NC licenses to GBIF (iNaturalist, 2025). In our case study, this excluded one of the two *Bombus impatiens* observations from Denton County, TX that is under copyright to the initial observer. It is possible to contact individual observers via the iNaturalist platform to obtain consent to incorporate copyrighted observations and obtain location information. However, the time and labor to facilitate permissions can be high and are not warranted for every project. Naturalist records obtained through GBIF also contain only the initial user who identified a record and not all users who contributed to the record. This may fail to identify observations that a taxon-specific expert has assessed and identified. Label photographs that have useful information for retrospective georeferencing are not directly visible via GBIF. Thus, for certain contexts a user may need to examine records through the original collection rather than relying solely on GBIF.

Caution must also be used with any “big data” approach. Large data streams from non-systematic sampling can produce biases (Boyd et al., 2023) that are important to consider when making inferences using such data. Biased inference can be mitigated by understanding the properties and limitations of specific data portals (Wüest et al., 2020), carefully cleaning data (Gueta and Carmel, 2016) using processes such as the workflow presented here, using formal risk-of-bias assessments or statistical adjustments (reviewed in Boyd et al., 2023), and validating models that use the data (Cazalis et al., 2022). Researchers, conservation practitioners, and policy makers should also carefully consider whether inaccuracies arising from “big data” approaches are relevant to the scale under consideration. Data may be robust to minor errors at the global scale but highly sensitive when looking at local patterns (e.g., Harry and Braccini, 2021). Alternately, a local survey effort may make knowledge of a species’ distribution high quality for that location but with biases across the species’ entire range (e.g., Rodríguez-Caro et al., 2024). The workflow described here, combined with R’s filtering options, has the flexibility to enable users to use the data that they feel are best for their goal and location and to implement the case-specific cleaning recommended by Gueta and Carmel (2016). When possible, management decisions should integrate information from both big data and experimental approaches to balance out the known errors associated with each approach (McCleery et al., 2023).

Future versions of our workflow, or customization by more advanced R users, could expand beyond current functionality. Making the workflow available through GitHub facilitates such expansion and improvement by the broader community. Taxon-specific cleaning functions, such as those for marine invertebrates (EcoCleanR R package, Soni et al., 2026) or fungi (Bates et al., 2025), can be added to our workflow. As discussed above, we do not include the option for automated georeferencing. As georeferencing tools become more reliable, for example through use of large language models (Fernando et al., 2026), automated georeferencing of records flagged as containing collection location descriptions could be added to our workflow. As we found with our case study, the Natural Earth maps that our third spatial filter uses can be rather inaccurate along coastlines, rivers, or other complex boundaries. Investment in improved maps would improve spatial filtering. Advanced R users could also write code to compare spatial filtering options and select the most reliable one. Another less challenging potential addition is quality control of user-entered dates and administrative divisions other than country. We did not automate date, stateProvince, or county harmonization due to lack of a global dataset with all possible values of these elements. However, code to standardize the values within a specific dataset is fairly trivial to write for a given geographic region and small dataset using, for example, the dplyr::replace_values() (Wickham et al., 2019) or by modifying BeeBDC::countryNameCleanR() to use a local maps file. A final addition that would increase utility of our workflow for NatureServe assessment is incorporating threat assessment that incorporates the scope, severity, and timing of specific anthropogenic or natural threats. While this can refine assessments, threat is difficult to quantify for many species with poorly understood natural histories. Overall, the workflow described here presents a significant advancement for aggregating, cleaning, and standardizing species occurrence records at scale in a transparent and flexible manner.

## Acknowledgements

We thank Victoria Vo for feedback. This project was funded US Fish and Wildlife Service Competitive State Wildlife Grants F21AF02917 and F22AF03320.

